# An acute session of motor imagery training induces use-dependent plasticity

**DOI:** 10.1101/716357

**Authors:** Célia Ruffino, Jérémie Gaveau, Charalambos Papaxanthis, Florent Lebon

## Abstract

Motor imagery, defined as the mental representation of an action without movement-related sensory inputs, is a well-known intervention to improve motor performance. In the current study, we tested whether use-dependent plasticity, a mechanism underlying motor learning, could be induced by an acute session of motor imagery. By means of transcranial magnetic stimulation (TMS) over the left primary motor cortex, we evoked isolated thumb movements in the right hand and assessed corticospinal excitability in the flexor and extensor pollicis brevis muscles. We measured the mean TMS-induced movement direction before and after an acute session of motor imagery practice. In a first experiment, participants of the imagery group were instructed to repeatedly imagine their thumb moving in a direction deviated by 90° from the pre-test movement. This group, but not the control group, deviated the post-training TMS-induced movements toward the training target direction (+44° ±62° and −1° ±23°, respectively). Interestingly, the deviation magnitude was driven by the corticospinal excitability increase in the agonist muscle. In a second experiment, we found that post-training TMS-induced movements were proportionally deviated toward the trained direction and returned to baseline 30 minutes after the motor imagery training. These findings suggest that motor imagery induces use-dependent plasticity and, this neural process is accompanied by corticospinal excitability increase in the agonist muscle.

## Introduction

Use-dependent plasticity is a basic neural mechanism underlying motor learning. At the behavioral level, consistently repeating reaching movements in a given direction has been shown to reduce movement variability and to bias future movements toward that direction ^1,2^. At the neural level, a well-known paradigm to test for the existence of use-dependent plasticity is to stimulate the primary motor cortex, by means of transcranial magnetic stimulations (TMS), before and after a motor training session, comprised of several repetitions of the same movement ^3–5^. If the direction of TMS-induced movement changes after the intervention, a use-dependent plasticity mechanism responsible for this bias could be inferred.

Previous studies have shown that physical practice ^3,5-7^, as well as action observation ^6,7^ induce use-dependent plasticity; i.e., post-intervention TMS-induced movements were deviated toward the trained direction. These effects were even potentiated when combining physical practice and action observation in aged individuals ^5^ and in stroke patients ^8^, thereby offering promising applications for neurorehabilitation. Physical practice and action observation both involve sensory feedback, either from one’s own movement during physical practice or from someone else’s movement during action observation. Whether use-dependent plasticity could be induced without movement-related sensory feedback, for instance by mentally repeating a movement, is still an open question.

Motor imagery is a promising intervention for neurorehabilitation ^9–11^. Classically defined as the mental representation of movement without its concomitant execution ^12^, motor imagery is an explicit process during which a participant is asked to recall the sensorimotor representations that are normally generated during actual execution. Although motor imagery has been long shown to improve motor performance ^13–15^, the underlying neural mechanisms are still largely unknown. Functionally, no movement-related sensory feedback is actually involved during motor imagery training, and, therefore, an error-based learning process ^16,17^ should be excluded. From a computational perspective, the activation of the corticospinal tract during repetitive imagined movements ^18^ could improve the function of the controller and, consequently, enhance motor performance ^19,20^. At a neural level, motor imagery training was considered to induce long-term potentiation-like plasticity ^21^. This neural modulation has been suggested to be associated with use-dependent plasticity following actual practice^4^. Therefore, use-dependent plasticity would be a good theoretical-based neural mechanism to explain motor improvements obtained with motor imagery training.

To probe the influence of an acute motor imagery session on use-dependent plasticity, we performed two experiments during which the participants were asked to imagine thumb movements at specific directions (motor imagery groups) or to stay at rest (control group). We first hypothesized that motor imagery would induce use-dependent plasticity, and that target movement deviation would be accompanied by modulation of corticospinal excitability (experiment 1). Then, we predicted that movement deviation would be proportional to the training direction with a short-lasting effect (experiment 2), as previously observed with physical practice ^3^.

## Materials and Methods

### Participants

Forty-five healthy volunteers, without neurological or physical disorders, participated in the current study after giving their informed consent. The study was approved by the CPP SOOM III ethics committee (ClinicalTrials.gov Identifier: NCT03334526) and complied with the standards set by the Declaration of Helsinki.

### Experimental device and procedure

In both experiments, the participants comfortably sat in a chair with their right dominant arm resting on a table in front of them. Their right forearm and hand were restrained in a brace, but thumb movements were unrestrained (Fig. 1.A). To evaluate use-dependent plasticity, we used TMS to induce artificial thumb movements at complete rest before (PreTest) and after (PostTest) motor imagery training.

**Figure 1.**
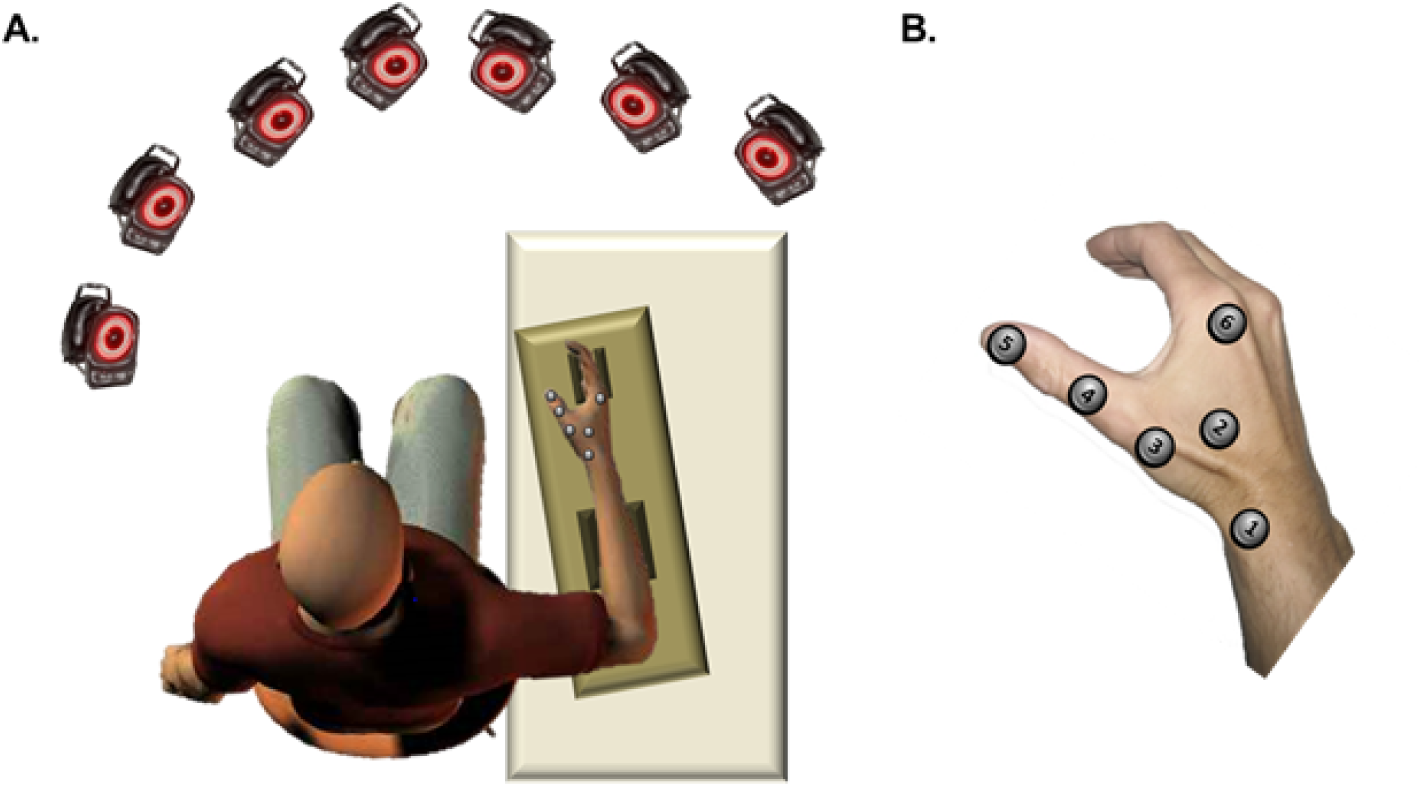
**A.** Position of the participant and the infra-red cameras. **B.** Position of retro-reflective markers.

In the PreTest session of both experiments, we delivered 60 TMS pulses at rest, the participants remained quiet and relax during the stimulations, and we recorded the direction of the elicited thumb movements (see Fig. 2.A). For each participant, the mean movement direction was computed immediately after the stimulations to define the subsequent direction for the motor imagery training. The PostTest was similar to PreTest, i.e., 60 TMS pulses were elicited and thumb movement direction was computed again. Finally, we tested the lasting effect of use-dependent plasticity by repeating the PostTest procedure 30 minutes (Post30) and 60 minutes (Post60) after the end of training, for one group only (the MI110 group, see below for group description).

**Figure 2.**
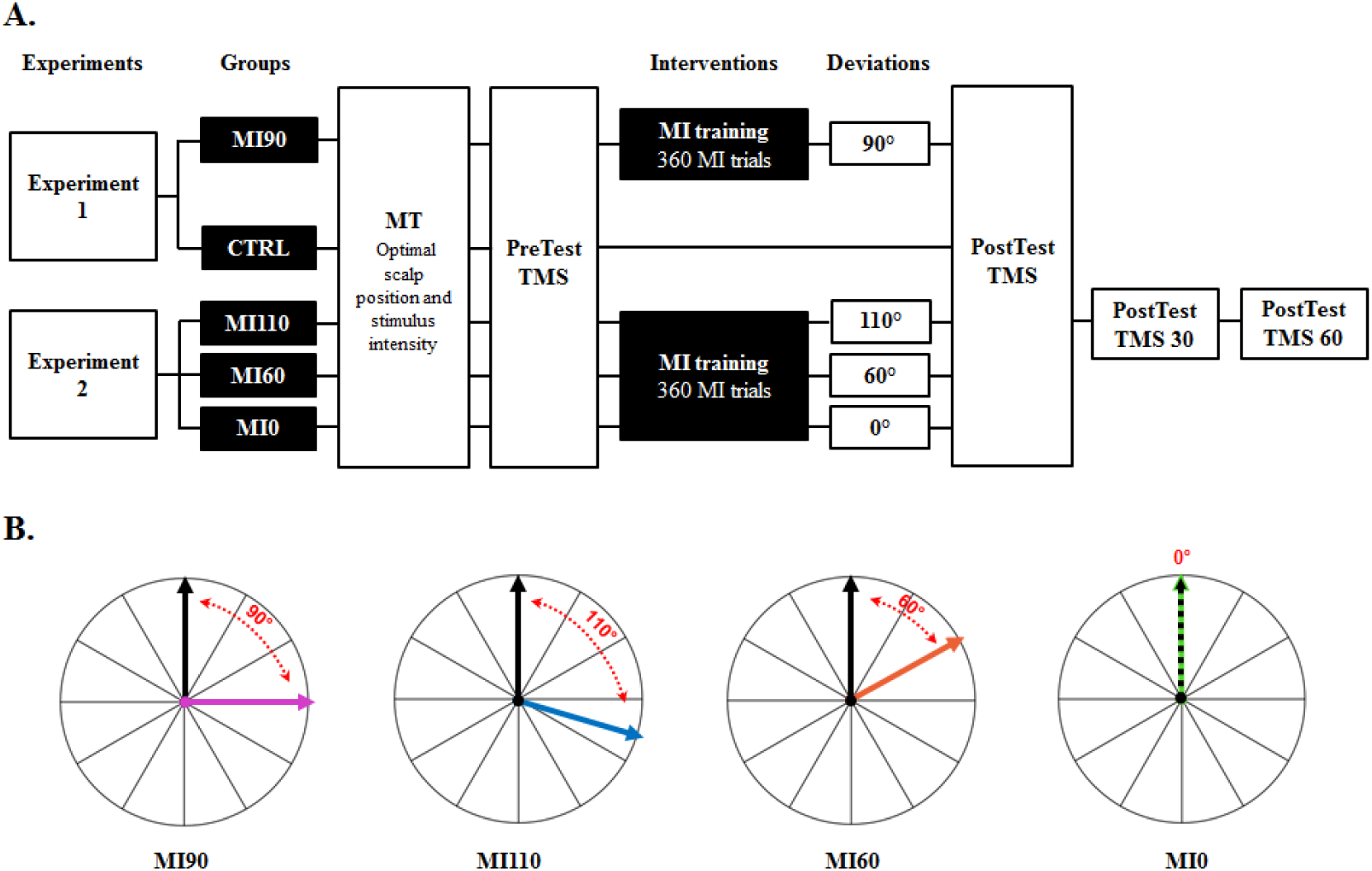
**A.** Intervention design. **B.** Illustration of the target angles used to train each group. The black arrow pictures the mean TMS-induced movement at PreTest and the colored arrow represents the direction of the imagined movements during training. Note that mean direction of PreTest TMS-induced movements varied across participants. MI = Motor Imagery; MT = Movement Threshold; TMS = Transcranial Magnetic Stimulation.

During motor imagery training, the participants of each group were instructed to imagine movements in a specific direction relative to that recorded in the PreTest (Fig. 2.B). Precisely, in Experiment 1, participants imagined movements that were deviated clockwise by +90° from the PreTest mean direction (MI90 group, n=12, 6 males, 27±8 years old). The control (CTRL) group of the Experiment 1 (n=12, 6 males, 31±11 years old) was instructed to not engage in any actual or mental repetition of thumb movement. Motor imagery ability of both groups (MI90 and CTRL) was assessed by the French version of the Movement imagery Questionnaire ‘MIQr’ ^22^. The average score was 44 ±6 and 45 ±6, respectively, indicating a good and equivalent imagery ability (t=0.03, P=0.97, Cohen’s d=0.17; maximum theoretical score: 56; minimum theoretical score: 8).

In Experiment 2, the first group (MI110, n=7, 5 males, 24±2 years old) imagined movements that were deviated clockwise by +110° from the PreTest mean direction, the second group (MI 60°, n=7, 6 males, 27±5 years old) imagined movements that were deviated clockwise by +60° from the PreTest mean direction, and the third group (MI0, n=7 males, 27±4 years old) mentally trained in the direction (0°) of the PreTest TMS-induced movement.

All motor imagery groups performed 360 imagined movements divided in 6 blocks of 60 trials with 1-min rest between blocks to avoid mental fatigue ^23^. The following instructions were provided: “try to imagine yourself performing the thumb movement in the trained direction, by feeling the finger sensation as if you were doing it and perceiving the movement direction just as if you were looking at your thumb moving into that direction”. The target with the trained direction illustrated by an arrow was placed in front of the participant to help imagining the proper direction.

### Transcranial Magnetic Stimulation

TMS pulses were delivered via a figure-of-eight-shaped coil (70-mm external wing diameter) attached to a Magstim 200 stimulator (Magstim Co, Whitland, Wales, UK). The center of the coil was positioned over the left primary motor cortex to evoke thumb movements in the right hand. The coil was held tangentially to the scalp, with the handle pointing backward and 45° away from the midline of the skull. The optimal scalp position was identified and the movement threshold was defined as the lowest intensity to evoke isolated thumb movements. The averaged stimulation intensity was 48 ±8% and 47 ±7% of maximum stimulator output in experiments 1 and 2, respectively.

Surface electromyography (EMG) was recorded from right extensor pollicis brevis (EPB) and flexor pollicis brevis (FPB) using bipolar electrode configurations. These muscles were chosen as they are major contributors to thumb extension and flexion. EMG was amplified (x1000), with a bandwidth frequency ranging from 10 Hz to 1 kHz, sampled at 2 kHz using a soft-ware commercially available (AcqKnowledge; Biopac Systems, Inc., Goleta, CA). During motor imagery training, absence of voluntary activation was monitored online by measuring EMG root mean square (EMGrms). If voluntary EMGrms activity was detected (i.e., 2.5 standard deviations around the mean EMGrms at rest), the participants were instructed to relax^8^. At Pre and PostTest sessions, EMGrms was also calculated 100 ms prior to TMS artefact to ensure that MEP amplitudes were not contaminated by muscle activation. Trials with EMGrms above 2.5 standard deviations from the mean within the same condition were removed for further analysis (3.1% of trials).

### Kinematics

Thumb movements were recorded at 200 Hz in all directions (X: mediolateral, Y: antero-posterior and Z: vertical) using a seven-camera motion capture system (Vicon, Oxford, UK). Six retro-reflective markers (diameter = 15 mm) were positioned on the skin at the following anatomical locations (see Fig.1.B): [1] Styloid process of the radius; [2] Middle of flexor pollicis brevis; [3] Metacarpal bone of thumb; [4] First phalanx of thumb [5] Second phalanx of thumb; [6] trapezoid. Using custom programs in Matlab (Mathworks, Natick, MA), the direction of each TMS-induced movement (in degree) was computed using classical vector geometry in three dimensions. Here, we report the thumb movement direction at the time of peak acceleration.

### Data analysis

#### Movement deviation

In previous studies, authors have mainly expressed use-dependent plasticity as the percentage of TMS-induced movements falling in a target zone defined around the trained direction. In the current study, we computed the mean TMS-induced movement direction (in degree). To do so, we first computed TMS-induced thumb displacements in 3D for each trial and then averaged each dimension across trials. The mean direction was finally computed from the averaged 3D deviations, therefore ensuring that the contribution of each trial was proportional to the amplitude of the TMS-induced thumb movement. To report inter-test thumb directional deviations in a similar frame of reference for all participants, we subtracted the PreTest mean direction to the PostTest one. We thus obtained normalized directional deviations, where 0° corresponds to no-deviation, whilst positive and negative values correspond to clockwise and counter-clockwise deviations, respectively.

#### Motor-evoked potential analysis

In Experiment 1, we determined whether a potential deviation toward the imagined movement direction was also accompanied by a modulation of corticospinal excitability. We measured motor-evoked potential (MEP) amplitude (in mV) at PreTest and PostTest in FPB and EPB muscles. To ascertain the contribution of the two muscles relative to the training direction (agonist vs. antagonist), we first determine the direction for full extension and full flexion when the wrist was positioned in the cast. Before the completion of the study, five participants were instructed to perform actual brisk movements toward extension and flexion and we measured the direction of movements as described above. In the current study, the extension/flexion axis was about 60° counterclockwise from the vertical axis, i.e., full extension was oriented up and to the left while full flexion was oriented down to the right (Fig. 3). From this analysis, we considered for each individual FPB as the agonist and EPB as the antagonist when the training target was toward full flexion, and conversely. For the agonist and antagonist muscles, we averaged MEP amplitudes for PreTest and PostTest sessions, removing trials when above 2.5 standard deviations from the mean (3.9% of trials). Finally, we calculated the ratio Post/Pre for the agonist and antagonist muscles in both MI90 and CTRL groups.

**Figure 3.**
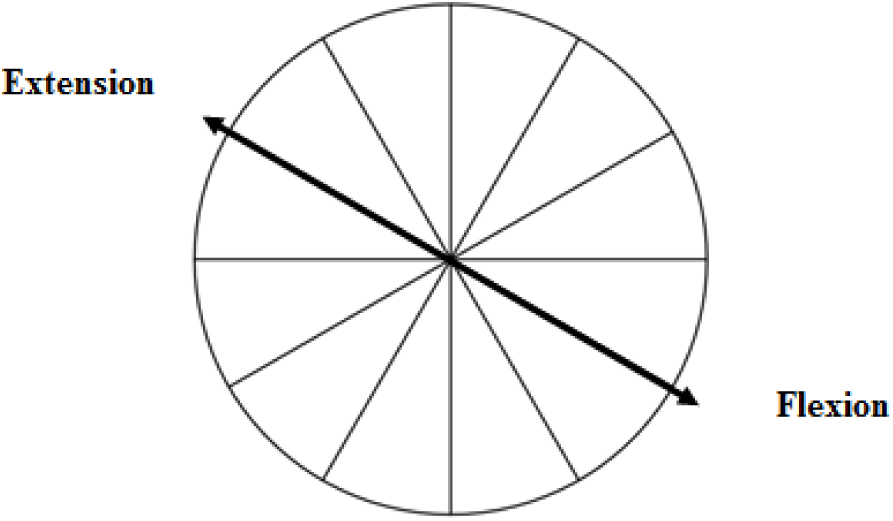
Illustration of actual brisk flexion and extension. When the arm and forearm were positioned in the cast, thumb flexion and extension were oriented about 60° counterclockwise from the vertical axis. This helped determining for each individual the role (agonist or antagonist) of flexor and extensor pollicis brevis muscles relative to the trained direction.

#### Statistical analysis

Normality of distribution was verified by the Shapiro-Wilk test. Homogeneity of variances was assessed by Mauchly’s test and Greenhouse-Geisser correction was applied if the sphericity assumption was violated. In both Experiments, we first compared normalized deviations to the reference value, i.e., 0, using one-sample t-tests. To compare the influence of imagined directions, we used a Student t-test (CTRL vs. MI90) in Experiment 1, and a Kruskal-Wallis rank-test (with Mann-Whitney tests for paired comparisons) with between-subject factor *Direction* (MI110, MI60, MI0) on normalized deviations in Experiment 2. To test the lasting effect of use-dependent plasticity following motor imagery, we compared normalized deviations between PostTests for the MI110 group in Experiment 2, with a Friedman test (and Wilcoxon tests for paired comparisons) with within-subject factor *Test* (Post0, Post30, Post60). To evaluate the modulation of corticospinal excitability in Experiment 1, we used a repeated-measure ANOVA with *Muscle* as within-subject factor (Agonist, Antagonist) and *Group* as between-subject factor (CTRL, MI90) on MEP amplitude ratios. One-sample t-tests were also used to compare normalized MEP amplitude to the reference value, i.e., 1. Finally, a multiple linear regression was calculated to predict participants’ deviation following motor imagery training toward 90° upon the MEP amplitude modulation between Pre- and Post-tests for agonist and antagonist muscles. To do so, normalized TMS-induced deviation was the dependent variable and Post/Pre MEP ratios of agonist and antagonist muscles were the independent variables, i.e., the predictors. To ensure that MEPs were not contaminated by muscle activation, we compared EMGrms between Pre and PostTests within the same condition, with Wilcoxon tests with correction for multiple comparisons.

## Results

### Experiment 1

First, we evaluate the effect of motor imagery training on TMS-induced thumb movement direction (see Fig.4). Movement direction was significantly deviated from PreTest in the MI90 group (+44 ±62°, CI_95%_=[4;83], t=2.44, P=0.03), but not in the CTRL group (−1±23°, CI_95%_=[-16;12], t=-0.27, P=0.79). The Student t-test showed a significant difference between the two groups (t=2.38, P=0.026; Cohen’s d=1.06). The deviation of the MI90 group was not correlated to MIQr score, i.e. individuals’ imagery ability (R=-0.05, P=0.88). These results suggest that motor imagery training can induce use-dependent plasticity.

**Figure 4.**
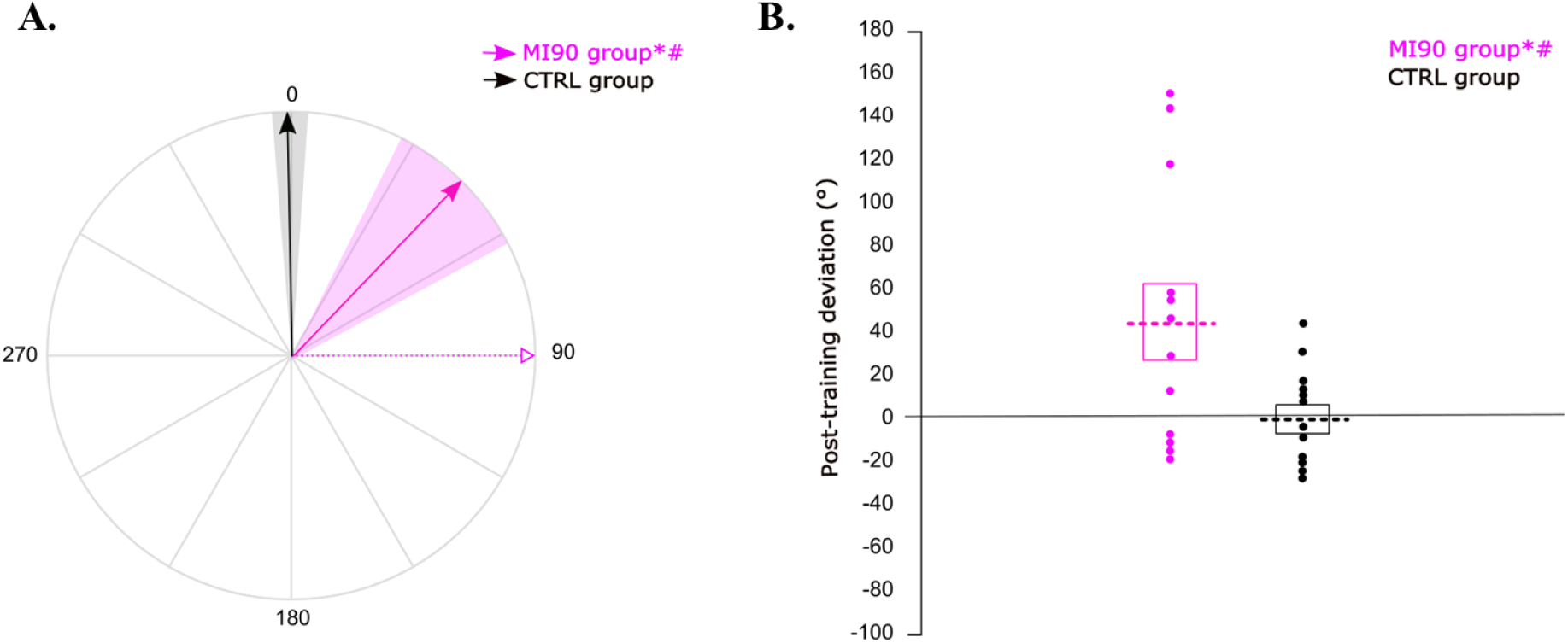
Post-training deviation. **A.** For each group, the solid arrow and the colored area correspond to the mean normalized deviation and to the standard error around the mean, respectively. The dotted pink arrow reminds the training direction for the motor imagery group. All participants are represented in a similar frame of reference, in which 0 is the TMS-induced movement direction at PreTest. **B.** For each group, the dotted lines correspond to the mean normalized deviation. The boxes represent the standard error of the mean and the dots refer to individual values. * = significantly different from PreTest (P<0.05); # = significant difference between MI90 and CTRL group (P<0.05).

We then test for the modulation of corticospinal excitability, comparing Post/Pre ratios of agonist and antagonist muscles between conditions. The rmANOVA did not reveal main effects (Muscle: F_1,22_=2.61, P=0.12, η_p_^2^=0.11; Group: F_1,22_=1.22, P=0.28, η_p_^2^=0.05) nor interaction (F_1,22_=2.72, P=0.11, η_p_^2^=0.11). Also, the ratios did not differ between Pre and Post sessions for both groups and muscles, even if the MI agonist ratio tends to increase at PostTest (One-sample t-tests, all P’s>0.1; MI agonist ratio = 1.63 ±1.37, CI_95%_=[0.76;2.50]; MI antagonist ratio = 1.07 ±0.59, CI_95%_=[0.69;1.45]; CTRL agonist ratio = 1.04 ±0.22, CI_95%_=[0.90;1.19]; CTRL antagonist ratio = 1.05 ±0.44, CI_95%_=[0.77;1.32]). This result corroborates the absence of corticospinal excitability increase after an acute session of motor imagery, classically observed in the literature ^21^.

While MEP amplitude increase did not reach significance after motor imagery training when tested as a group effect, we questioned whether individual MEP amplitude in agonist and antagonist muscles could predict movement deviation using a multiple linear regression. Interestingly, we found a significant regression equation (F_2,9_=8.20, p=0.009), with an adjusted R^2^ of 0.57. The movement deviation following motor imagery training toward 90° was predicted by the agonist ratio (Beta=0.84, partial correlation=.74, P=0.009, Figure 5), but not the antagonist ratio (Beta=-0.06, partial correlation=-.08, P=0.81). Tests to verify whether data met the assumption of collinearity indicated that multicollinearity was not a concern (agonist ratio, Tolerance = .59, VIF = 1.69; antagonist ratio, Tolerance = .59, VIF = 1.69). This finding indicates that the greater the increase in agonist MEPs between Pre and PostTests, the greater the deviation following motor imagery training. Figure 6 shows a typical representation of TMS-induced movement direction and MEP amplitude modulation in two participants.

**Figure 5.**
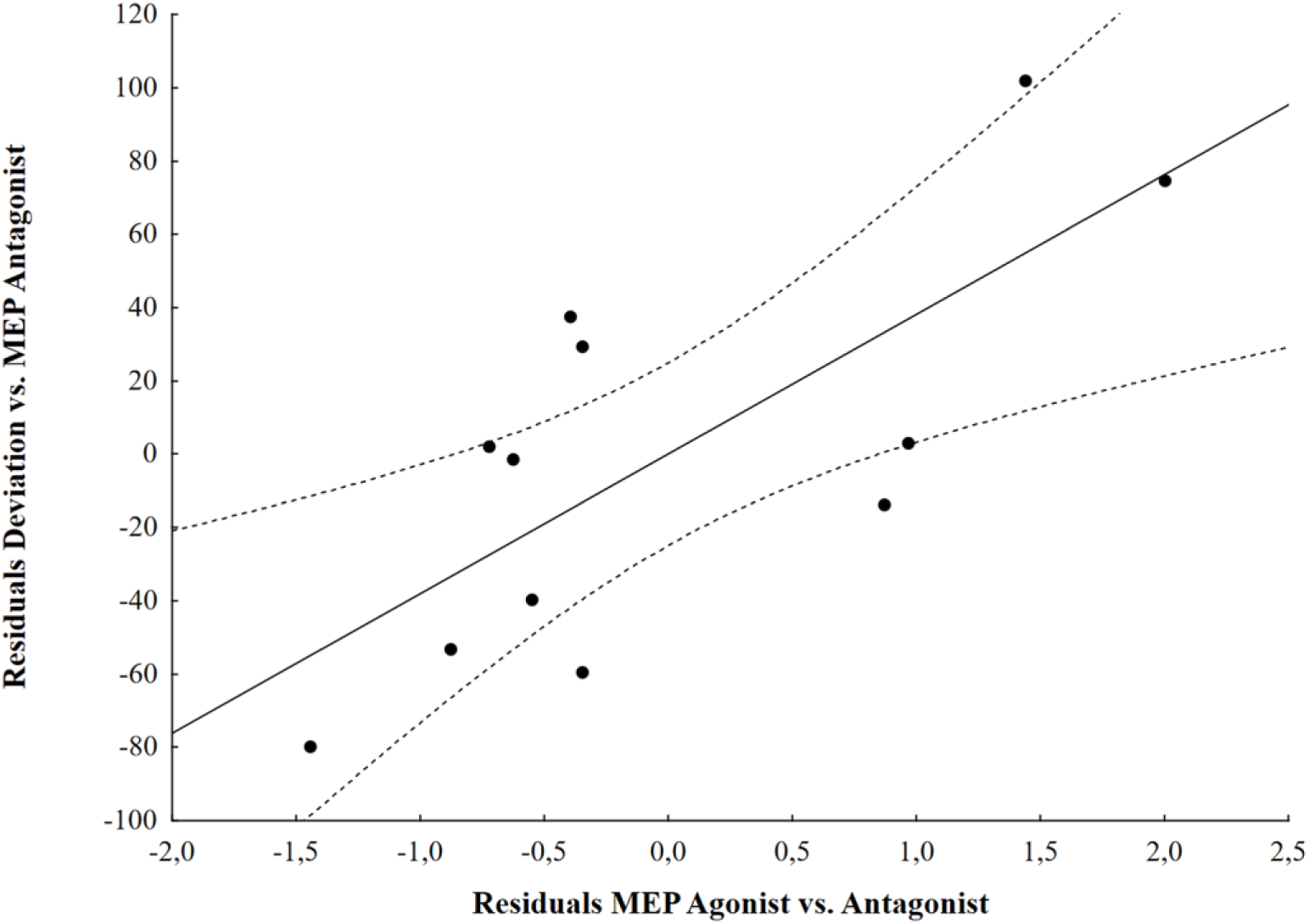
Partial scatter plot of the association between the movement deviation and the ratio MEP Agonist when controlling for the ratio MEP Antagonist (partial correlation=.74, P=0.009). Solid line = fitting regression line; dotted line = 95% confidence regression bands.

**Figure 6.**
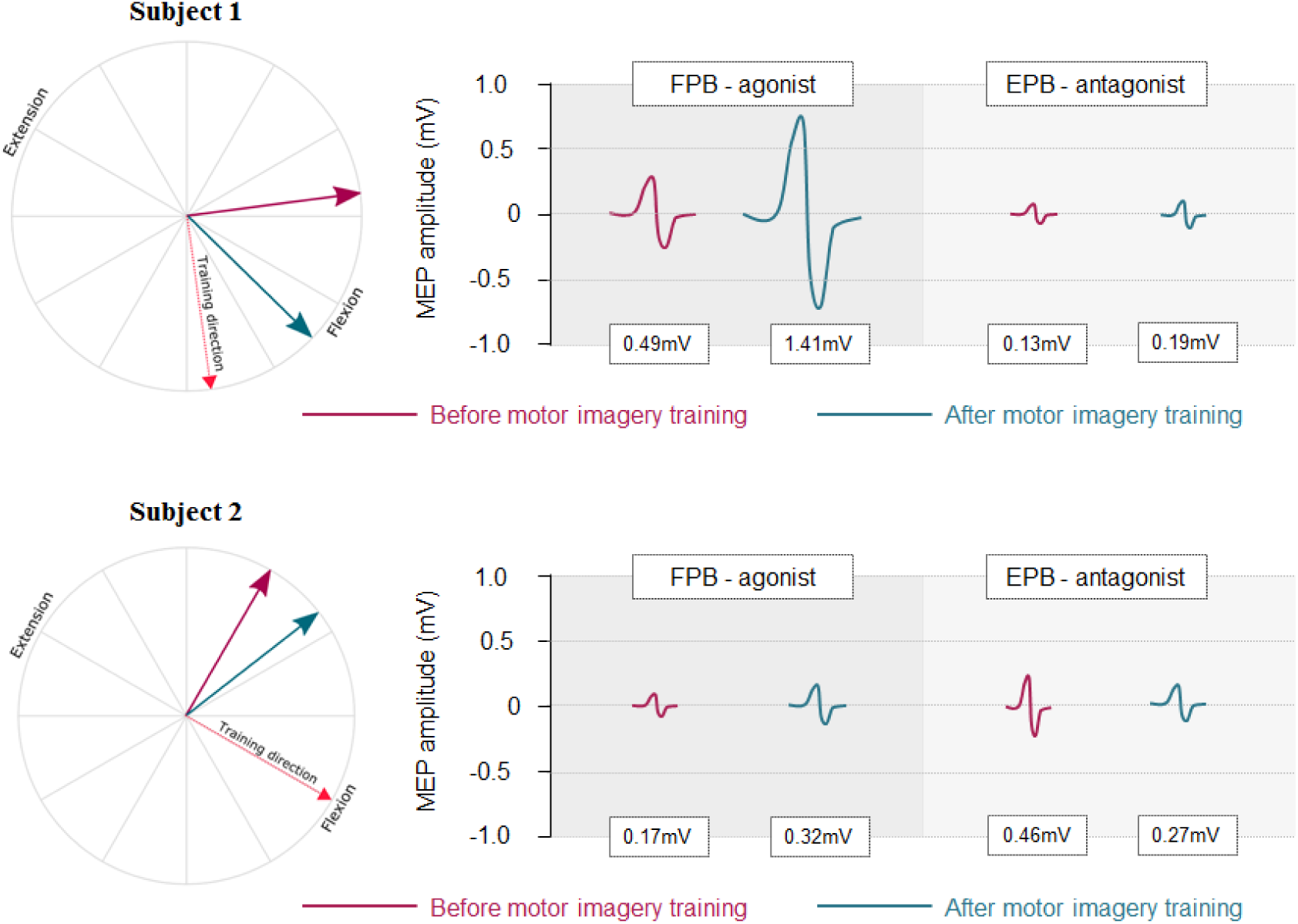
Typical representation of TMS-induced movement direction and MEP amplitude in flexor pollicis brevis (FPB) and extensor pollicis brevis (EPB) muscles in two subjects of the MI90 group. In both subjects, FPB was the agonist of the trained movement. Subject 1 deviated from Pre to PostTest by +52° and increased MEP amplitude in the agonist and antagonist muscles by 215% and 51%, respectively. Subject 2 deviated from Pre to PostTest by +22° and increased and decreased MEP amplitude in the agonist and antagonist muscles by 88% and 42%, respectively.

EMGrms 100 ms prior to TMS artefact did not differ between conditions (in all, P’s>0.25; means ±SD for MI90 group: PreTest FPB = 3.25 ±4.32 µV, PostTest FPB = 3.93 ±4.53 µV, PreTest EPB 2.90 ±1.31 µV, PostTest EPB = 3.46 ±1.97 µV; means ±SD for CTRL group: PreTest FPB = 5.72 ± 6.05 µV, PostTest FPB = 5.80 ±5.16 µV, PreTest EPB = 5.56±4.05 µV, PostTest EPB = 6.18 ±6.03 µV). This ensured that MEP amplitudes and TMS-induced deviations were not contaminated by muscle activity.

During motor imagery training, mean EMGrms during imagery trials and at rest were 2.70 ±0.89µV and 2.10 ±0.81 µV for FPB and 4.63 ±2.36 µV and 3.35 ±1.51 µV for EPB, respectively. The difference between the conditions is not significant (all P’s>.10).

### Experiment 2

In a second experiment, we evaluated the specific effect of motor imagery training on movement deviation and the lasting effect of use-dependent plasticity. Figure 7A illustrates the effect of target angle on the normalized directional deviation. After motor imagery training, TMS-induced movements were significantly deviated from PreTest in the MI60 group (+40 ±30°, CI_95%_=[11;68], t=3.46, P=0.01) and in the MI110 group (+89 ±51°, CI_95%_=[41;137], t=4.57; P=0.003), but not in the MI0 group (−7 ±10°, CI_95%_=[-16;3], t=-1.66, P=0.15). Strikingly, the amplitude of the deviation increased with target angle, as testified by the significant effect of the trained angle (H=10.69, P=0.005, η^2^H=0.48; post-hoc MI110 vs MI0, Z=3.13, P<0.001, r^2^=0.70; post-hoc MI110 vs MI60, Z=1.72, P=0.08, r^2^=0.21; post-hoc MI60 vs MI0, Z=2.49, P=0.01, r^2^=0.43). We also found a positive correlation measured between the target angle and the normalized deviation (R-Spearman=0.79, p<0.001).

**Figure 7.**
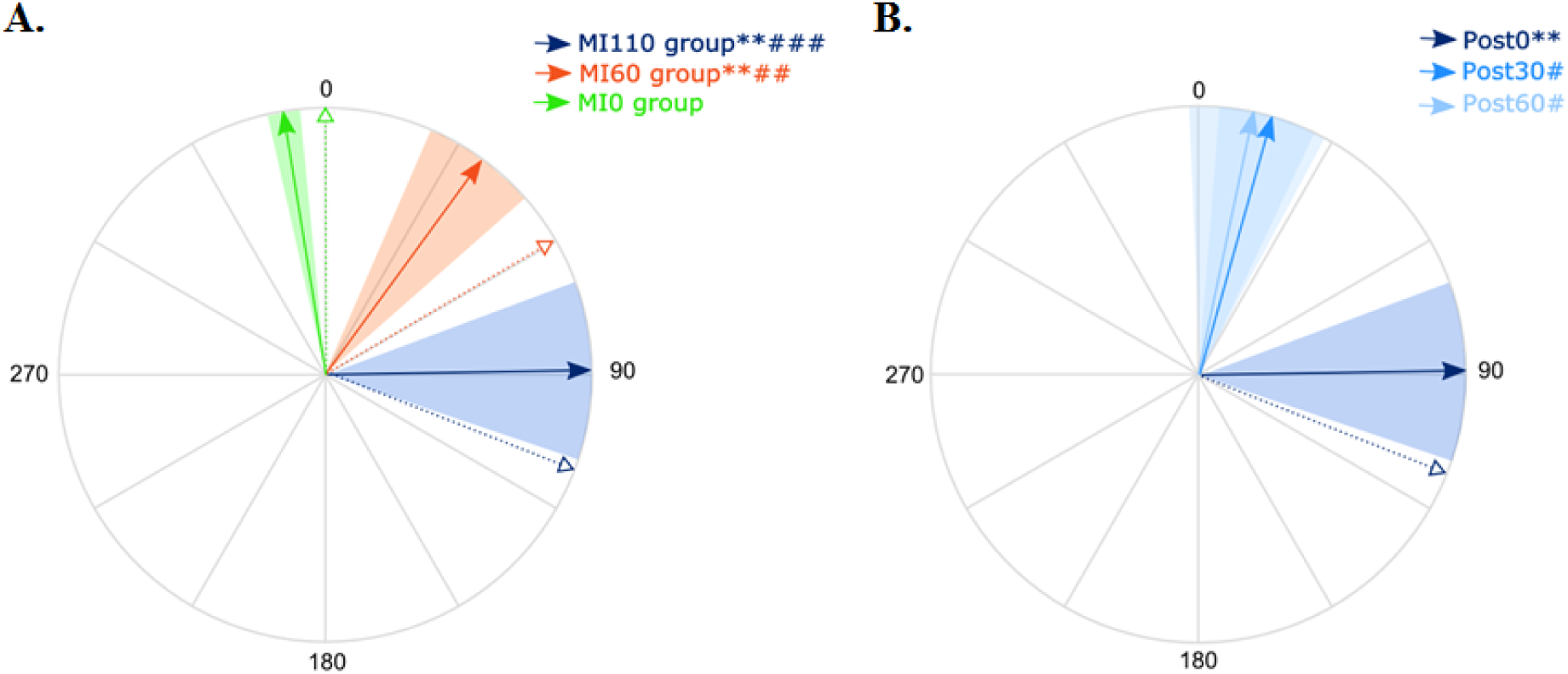
**A.** Post-training deviation. For each group, the solid arrow and the colored area correspond to the mean normalized deviation and to the standard error around the mean, respectively. The dotted arrow reminds the training direction for each group. All participants are represented in a similar frame of reference, in which 0 is the TMS-induced movement direction at PreTest. ** = significantly different from PreTest for MI60 and MI110 (P=0.01 and P=0.003 respectively); ## = significant different between MI60 and MI0 (P=0.01) and ### = significant difference between MI110 and MI0 (P<0.001). **B.** Lasting effects of use-dependent plasticity for the MI110 group. * and ** = significantly different from PreTest (P<0.05 and P<0.01, respectively); # and ### = significantly different from MI110 Post-test0 (P<0.05 and P<0.001, respectively).

Next, we test the lasting effects of motor imagery on use-dependent plasticity. Figure 7B reveals that the deviation observed just after motor imagery training for the MI110 group (+89 ±51°) attenuated with time, as testified by the significant effect of *Test* (χ^2^=8.0; P=0.018, W=0.19) and significant differences between Post0 and Post30 (Z=2.19, P=0.028, r^2^=0.34) and between Post0 and Post60 (Z=2.36, P=0.018, r^2^=0.40). The deviation was no longer significant in comparison to PreTest at Post30 (+16° ±31, CI_95%_=[-13;45], t=1.33, P=0.23) nor at Post60 (+12° ±36, CI_95%_=[-22;45], t=0.85, P=0.43).

## Discussion

In the current study, use-dependent plasticity was induced by an acute session of motor imagery practice. At the neurophysiological level, this process was merely driven by a MEP increase in the agonist muscle. Behaviorally, it was direction-specific and short-lasting, i.e., post-training TMS-induced movements were proportionally deviated toward the trained direction and returned to baseline when tested after 30 minutes. Our results closely match those previously reported following physical practice, as Classen et al. (1998) ^3^ in their seminal study also observed such a direction specific and short-lasting effect (see Figure 2.B and 2.C in Classen et al., 1998 ^3^).

An increase of corticospinal excitability associated with motor imagery training may explain the induction of use-dependent plasticity. Several studies have shown an increase in the amplitude of MEPs, a marker of corticospinal excitability, during motor imagery in a time- and muscle-specific manner ^18-20^. Interestingly, Sommer et al. (2001) ^24^ found that TMS-induced movements were deviated toward the intended direction from 90 ms prior to voluntary initiation. At that time point, MEP amplitude already increased in the target muscle. They hypothesized that a certain level of facilitation of the predominant muscle is necessary to generate kinematic changes. Use-dependent plasticity could originate from this muscle-specific facilitation. Our findings, especially the results of the multiple regression analysis, corroborate this hypothesis. Indeed, we found that the TMS-induced movement deviation was driven by the post-imagery increase in MEP amplitude of the agonist muscle. As an example, when the participant imagined thumb flexion during training, the amount of movement deviation post-training was predicted by the amount of excitability increase in the FPB muscle between Pre and Posttests. This is in line with the study by Ranganathan et al. (2004) ^25^ during which a 12-week motor imagery training induced strength increase of the elbow flexors with increase in EMG activity in the agonist muscle (i.e., biceps brachii) but not in the antagonist one (i.e., triceps brachii). The agonist/antagonist ratio was therefore in favor of the agonist muscle.

One study may challenge this hypothesis of muscle-specific facilitation. Meintzschel and Ziemann (2006) ^26^ proposed that the increase in corticospinal excitability of the agonist muscle observed with physical practice is neither sufficient (see also Flöel et al. 2005 ^27^) nor necessary to explain TMS-induced thumb movement deviations. The authors observed that an injection of cabergoline, a precursor of dopamine, reduced MEP amplitude but induced use-dependent plasticity. However, as raised by the authors themselves, one explanation could be that a strong decrease of antagonist muscles excitability lead to an improved signal-to-noise ratio in favor of the training agonist. The gain between agonist and antagonist muscles may be of importance to observe use-dependent plasticity.

It is now well accepted that the neurophysiological processes underlying use-dependent plasticity occur at the cortical level. Interestingly, studies in animals have revealed that the dynamic force (or the change in force) and the direction of intended movements can be observed within M1^24^. The direction of the population vector, computed from several individual neurons in monkey brains, was shown to change toward the direction of a visual target during the reaction time of the actual movement ^28^. Accordingly, the repeated activation of the motor system during motor imagery practice may reinforce a network as the result of Hebbian changes in the motor cortex ^2,29–32^. As for physical practice, long-term potentiation (LTP)-like plasticity may be a good candidate to induce such Hebbian plasticity following motor imagery ^33^. Avanzino et al. (2015) ^21^ have described such a mechanism after motor imagery training, using the paired associative stimulation technique. They found a reversal effect of LTP-like plasticity, i.e. reduced corticospinal excitability, following paired associative stimulation when preceded by motor imagery practice, whilst increased corticospinal excitability is classically observed after paired associative stimulation alone.

Could use-dependent plasticity by motor imagery practice occur at the spinal level as well? Recent evidence revealed the presence of a subliminal voluntary drive going along the corticospinal tract during motor imagery ^34^. This voluntary drive reaches the spinal level without activating alpha-motoneurons but modulates the excitability of spinal low-threshold pre-synaptic interneurons. The repetition of imagined movements has been hypothesized to increase the excitability at the Ia afferent-motoneuron junction by decreasing the basal inhibitory level ^18^. However, whilst action observation also induces use-dependent plasticity ^6^, it was shown to induce reversed spinal modulations in comparison to motor imagery and actual execution. The amplitude of H-reflex in finger flexor muscles, agonist of the action, decreased during observation of hand closing, and vice versa ^35^. Furthermore, the metabolic activity in the spinal cord of monkeys observing reach-to-grasp movements was reduced, likely to suppress the descending motor output ^36^. While it seems unlikely that use-dependent plasticity takes place at the spinal level after motor imagery training, further studies may challenge this hypothesis.

In conclusion, the current study revealed that an acute session of motor imagery practice induced use-dependent plasticity and this phenomenon is driven by an increase of corticospinal excitability in the agonist muscle. This finding expands previous behavioral studies that showed motor performance improvement after mental training and unmasks the underlying neural mechanisms. The key implication of this study considers that the presence of movement-related sensory feedback, as during actual execution and action observation, is not a prerequisite to induce such plasticity. In this way, modulating neural and behavioral components of the movement with motor imagery training may have high potential for motor rehabilitation.

## Acknowledgement

This work was supported by the French “Investissements d’Avenir” program, project ISITE-BFC (contract ANR-15-IDEX-0003). We thank Carol Madden-Lombardi for proofreading the manuscript.

## Author contribution

FL, CP, JG designed the experiments; CR collected the data; FL, CP, JG, CR analyzed the data; FL, CP performed statistics; FL, CP, JG, CR wrote the manuscript

